## Supplemental Figures and Tables for "The *C. elegans* TspanC8 tetraspanin TSP-14 exhibits isoform-specific localization and function"

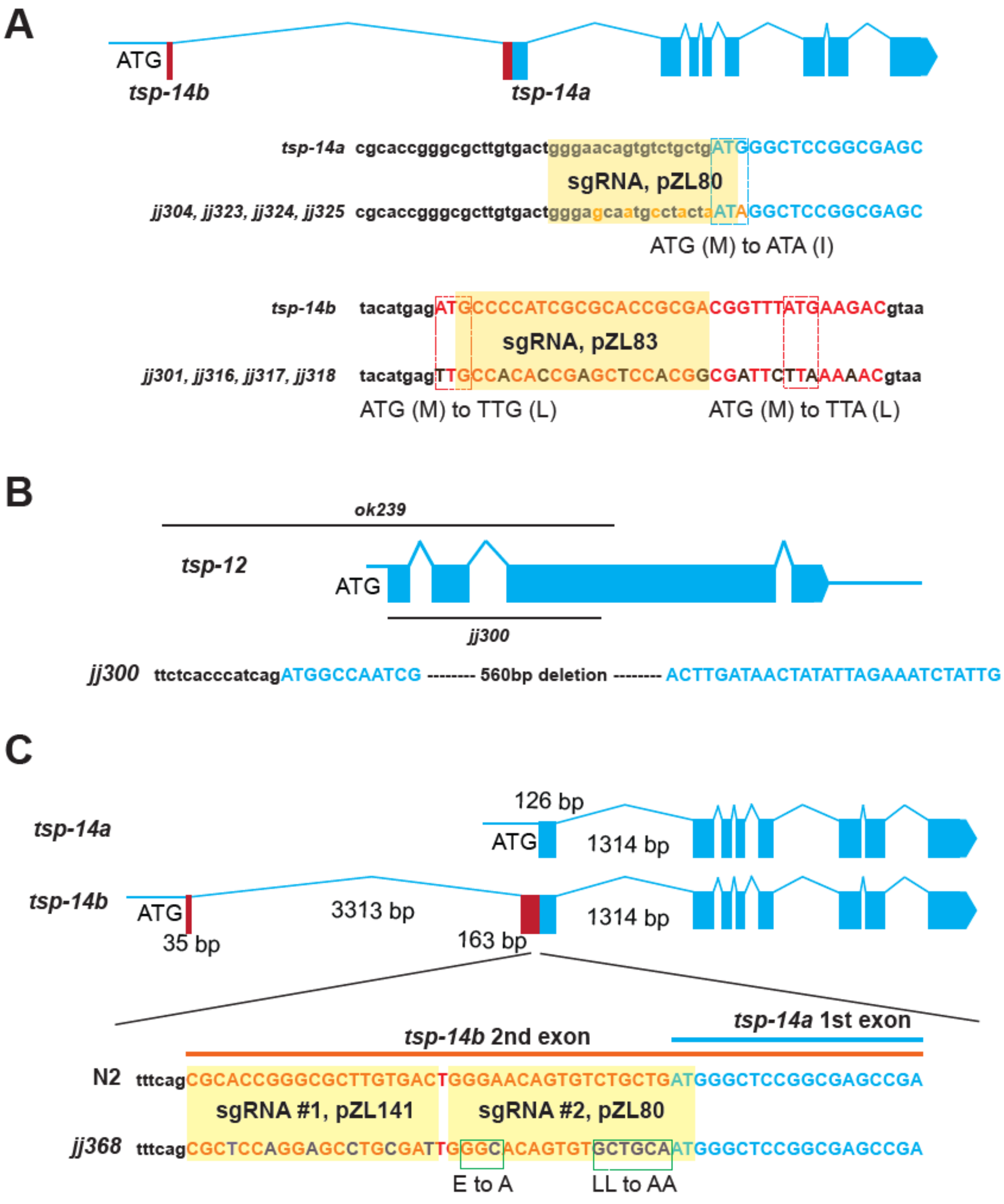
A
ATG
tsp-14b
tsp-14a
tsp-14a cgcaccgggcgcttgtgactgggaacagtgtctgctgATGGGCTCCGGCGAGC
sgRNA, pZL80
jj304, jj323, jj324, jj325 cgcaccgggcgcttgtgactgggagcaatgcctactaATAGGCTCCGGCGAGC
ATG (M) to ATA (I)
tsp-14b tacatgagATGCCCCATCGCGCACCGCGACGGTTTATGAAGACgtaa
sgRNA, pZL83
jj301, jj316, jj317, jj318 tacatgagTTGCCACACCGAGCTCCACGGCGATTCTTAAAAACgtaa
ATG (M) to TTG (L)
ATG (M) to TTA (L)
B
ok239
tsp-12
ATG
jj300
jj300 ttctcacccatcagATGGCCAATCG -------- 560bp deletion -------- ACTTGATAACTATATTAGAAATCTATTG
C
tsp-14a
126 bp
ATG
1314 bp
tsp-14b
ATG
35 bp
3313 bp
163 bp
1314 bp
tsp-14a 1st exon
tsp-14b 2nd exon
N2 tttcagCGCACCGGGCGCTTGTGACTGGGAACAGTGTCTGCTGATGGGCTCCGGCGAGCCGA
sgRNA #1, pZL141
sgRNA #2, pZL80
jj368 tttcagCGCTCCAGGAGCCTGCGATTGGGCACAGTGTGCTGCAATGGGCTCCGGCGAGCCGA
E to A
LL to AA

**S1 Figure**

**S1 Figure. Information on the different mutations in *tsp-12* or *tsp-14* generated using CRISPR/Cas9. (A)** *tsp-14* isoform specific knockout mutations. To specifically knock out *tsp-14a*, the start codon ATG (Met) of TSP-14A was mutated to ATA (Ile), to minimize significant impact on TSP-14B. To specifically knock out *tsp-14b*, the start codon ATG and another downstream, in-frame, ATG were mutated to TTG (Leu) and TTA (Leu), respectively. The sgRNA target site, as well as silent substitutions introduced to avoid cleavage of the repair templates by Cas9, are highlighted. **(B)** Schematics depicting the location of the two null mutations in *tsp-12*, *ok239* and *jj300*. The *jj300* allele was generated using CRISPR/Cas9, and the sequence of *jj300* is shown. **(C)** The EQCLL to AQCAA mutation in *tsp-14b*. Two sgRNAs used in this experiment. The sgRNA target sites, as well as silent substitutions introduced to avoid cleavage of the repair templates by Cas9, are highlighted. In all panels, coding sequences are in capital letters, while non-coding sequences are in lower cases.

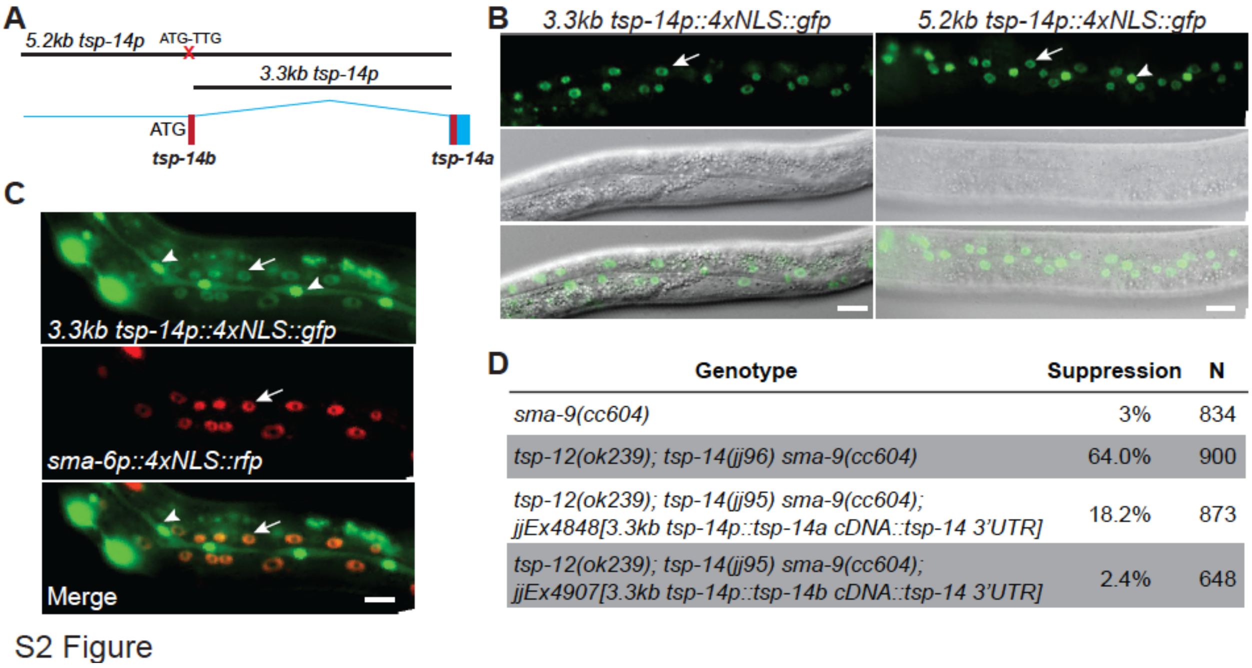

| Genotype | Suppression | N |
|---|---|---|
| *sma-9(cc604)* | 3% | 834 |
| *tsp-12(ok239); tsp-14(jj96) sma-9(cc604)* | 64.0% | 900 |
| *tsp-12(ok239); tsp-14(jj95) sma-9(cc604); jjEx4848[3.3kb tsp-14p::tsp-14a cDNA::tsp-14 3'UTR]* | 18.2% | 873 |
| *tsp-12(ok239); tsp-14(jj95) sma-9(cc604); jjEx4907[3.3kb tsp-14p::tsp-14b cDNA::tsp-14 3'UTR]* | 2.4% | 648 |

S2 Figure

**S2 Figure. Transgenic reporters of *tsp-14*. (A)** Schematics of the two different *tsp-14* promoters used. They are 3.3kb and 5.2kb, respectively, upstream of the *tsp-14a* start codon. In constructs with the 5.2kb promoter, the start codon of *tsp-14b* was changed from ATG to TTG. **(B)** Images showing GFP expression in hypodermal cells in transgenic animals carrying either of the two *tsp-14* transcriptional reporters. Scale bar, 20 µm. **(C)** Images showing that the 3.3kb *tsp-14* promoter drives GFP reporter expression in hypodermal cells that also express a *sma-6p:4xNLS::RFP* reporter. Notably, the 3.3kb *tsp-14p::4xNLS::GFP,* but not the *sma-6p:4xNLS::RFP* reporter, is expressed in the seam cells. Arrows point to hypodermal cell nuclei. Arrowheads point to seam cell nucleus. Scale bar, 20 µm. (D) Table summarizing the penetrance of the Susm phenotype of *tsp-12(0); tsp-14(0)* animals with or without the transgene expressing *tsp-14a* or *tsp-14b*. *tsp-14(jj95)* and *tsp-14(jj96)* are both null alleles of *tsp-14* and behave identically, as reported previously (Wang et al., 2017).

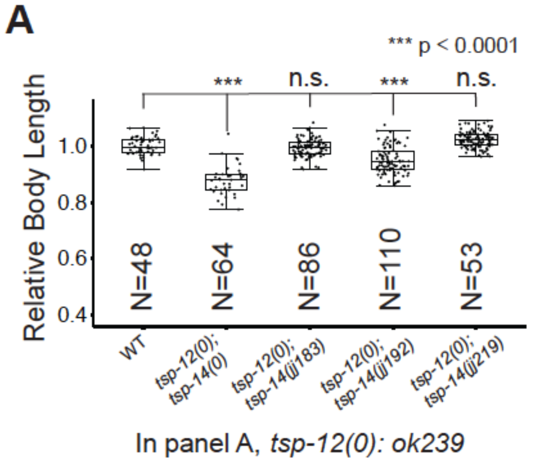

**B**

| Genotype | Suppression(%) | N |
|---|---|---|
| *sma-9(cc604)* | 1.5±0.4* | 476 |
| *tsp-12(ok239); sma-9(cc604)* | 13.1±1.8 | 343 |
| *tsp-14(jj96) sma-9(cc604)* | 1.5±0.4 | 502 |
| *tsp-12(ok239); tsp-14(jj96) sma-9(cc604)* | 64.0±1.0 | 505 |
| *tsp-14(jj183) sma-9(cc604)* | 0.9±0.4 | 616 |
| *tsp-12(ok239); tsp-14(jj183) sma-9(cc604)* | 36.0±0.9 | 718 |
| *tsp-14(jj192) sma-9(cc604)* | 1.7±0.4 | 424 |
| *tsp-12(ok239); tsp-14(jj192) sma-9(cc604)* | 6.9±0.3 | 696 |

S3 Figure

**S3 Figure. The functionality of *tsp-14* knock-ins in the *tsp-12(ok239)* background. (A)** Relative body lengths of synchronized L4 worms of various genotypes. The mean body length of wild-type worms is normalized to 1.0. Numbers within each bar indicate the total number of worms measured. For each double mutant, data were pooled from two independent isolates. Error bars represent 95% C.I. Tukey's HSD test following an ANOVA was used to test for differences between different genotypes. ***$P < 0.0001$. n.s., no significant difference. As shown, *tsp-12(0); tsp-14(jj192)* worms are slightly smaller than wild-type (WT) worms, but not as small as *tsp-12(0); tsp-14(0)* worms. **(B)** Table summarizing the results of the *sma-9(0)* suppression assay of various *tsp-14* knock-in alleles, in combination *tsp-12(ok239)*. Percentage of suppression was calculated by the number of worms with 1-2 M-derived CCs divided by the total number of worms scored. N represents the total number of worms counted. Data from two independent isolates were combined for each genotype. * The loss of M-derived CCs phenotype of *sma-9(cc604)* null mutants is not fully penetrant.

**Table S1. Strains genereated in this study.**

| Genotype | Strain Number |
|---|---|
| *tsp-12* and *tsp-14* null mutations | |
| *tsp-12(ok239)* | LW3382 |
| *tsp-12(jj300)* | LW5809 |
| *tsp-14(jj95)* | LW3713 |
| *tsp-14(jj96)* | LW3714 |
| *tsp-14a* isoform specific knockout | |
| *tsp-14(jj304: tsp-14a(ATG-ATA))* | LW5836 |
| *tsp-14(jj323: tsp-14a(ATG-ATA))* | LW5837 |
| *tsp-14(jj324: tsp-14a(ATG-ATA))* | LW5838 |
| *tsp-14(jj325: tsp-14a(ATG-ATA))* | LW5839 |
| *tsp-14b* isoform specific knockout | |
| *tsp-14(jj301: tsp-14b(ATG-TTG & ATG-TTA))* | LW5787 |
| *tsp-14(jj316: tsp-14b(ATG-TTG & ATG-TTA))* | LW5866 |
| *tsp-14(jj317: tsp-14b(ATG-TTG & ATG-TTA))* | LW5867 |
| *tsp-14(jj318: tsp-14b(ATG-TTG & ATG-TTA))* | LW5868 |
| TSP-14B sorting signal mutation | |
| *tsp-14(jj368: tsp-14b(EQCLL-AQCAA))* | LW5998 |
| *tsp-14(jj192 jj368: gfp::3xFLAG::tsp-14b(EQCLL-AQCAA))* | LW5743 |
| *tsp-14(jj368 jj378: tsp-14b(EQCLL-AQCAA)::gfp::3xFLAG)* | LW6043 |
| Endogeniously tagged proteins | |
| *tsp-12(jj181: gfp::3xFLAG::tsp-12)* | LW4453 |
| *tsp-14(jj219: tsp-14::gfp::3xFLAG)* | LW4768 |
| *tsp-14(jj183: gfp::3xFLAG::tsp-14a)* | LW4455 |
| *tsp-14(jj184: gfp::3xFLAG::tsp-14a)* | LW4456 |
| *tsp-14(jj186: gfp::3xFLAG::tsp-14a)* | LW4486 |
| *tsp-14(jj200: TagRFP::3xMyc::tsp-14a)* | LW4574 |
| *tsp-14(jj201: TagRFP::3xMyc::tsp-14a)* | LW4575 |
| *tsp-14(jj192: gfp::3xFLAG::tsp-14b)* | LW4520 |
| *tsp-14(jj193: gfp::3xFLAG::tsp-14b)* | LW4521 |
| *tsp-14(jj202: TagRFP::3xMyc::tsp-14b)* | LW4566 |
| *tsp-14(jj326: TagRFP::3xMyc::tsp-14b)* | LW4576 |
| *tsp-14(jj304 jj319: tsp-14::GFP::3xFLAG)* | LW5895 |
| *tsp-14(jj304 jj327: tsp-14::GFP::3xFLAG)* | LW5896 |
| *tsp-14(jj317 jj377: tsp-14::GFP::3xFLAG)* | LW6023 |
| Knock-in functionality | |
| *gfp::3xFLAG::tsp-14a(jj183) sma-9(cc604); ccIs4438* | |
| *gfp::3xFLAG::tsp-14b(jj192) sma-9(cc604); ccIs4438* | LW6067 |
| *tsp-12(ok239); gfp::3xFLAG::tsp-14a(jj183)* | LW4506, LW4507 |
| *tsp-12(ok239); gfp::3xFLAG::tsp-14b(jj192)* | LW4598, LW4599 |
| *tsp-12(ok239); gfp::3xFLAG::tsp-14a(jj183) sma-9(cc604); ccIs4438* | LW5219, LW5220 |
| *tsp-12(ok239); gfp::3xFLAG::tsp-14b(jj192) sma-9(cc604); ccIs4438* | LW5211 |
| *tsp-12(jj300); gfp::3xFLAG::tsp-14a(jj183)* | LW6031, LW6032 |

| | |
|---|---|
| *tsp-12(jj300); gfp::3xFLAG::tsp-14b(jj192)* | LW6037, LW6044 |
| *tsp-12(jj300); gfp::3xFLAG::tsp-14a(jj183) sma-9(cc604); ccIs4438* | LW6038, LW6039 |
| *tsp-12(jj300); gfp::3xFLAG::tsp-14b(jj192) sma-9(cc604); ccIs4438* | LW6063, LW6065 |
| Knock-out functionality | |
| *nT1[qIs51]/tsp-12(jj300); tsp-14(jj95)* | LW5830, LW5831 |
| *nT1[qIs51]/tsp-12(jj300); tsp-14(jj304: tsp-14a(ATG-ATA))* | LW5873 |
| *tsp-12(jj300); tsp-14(jj317: tsp-14b(ATG-TTG & ATG-TTA))* | LW5950, LW5951 |
| *tsp-12(jj300); tsp-14(jj368: tsp-14b(EQCLL-AQCAA))* | LW6006, LW6007 |
| *sma-9(cc604); ccIs4438* | LW3711 |
| *tsp-12(jj300); ccIs4438* | LW5991, LW5990 |
| *tsp-14(jj95) sma-9(cc604); ccIs4438* | LW6021, LW6022 |
| *nT1[qIs51]/tsp-12(jj300); tsp-14(jj95) sma-9(cc604); ccIs4438* | LW6048 |
| *tsp-14(jj304: tsp-14a(ATG-ATA)) sma-9(cc604); ccIs4438* | LW5874, LW5875 |
| *nT1[qIs51]/tsp-12(jj300); tsp-14(jj304: tsp-14a(ATG-ATA)) sma-9(cc604); ccIs4438* | LW5900, LW5901 |
| *tsp-14(jj317: tsp-14b(ATG-TTG & ATG-TTA)) sma-9(cc604); ccIs4438* | LW5948, LW5926 |
| *tsp-12(jj300); tsp-14(jj317: tsp-14b(ATG-TTG & ATG-TTA)) sma-9(cc604); ccIs4438* | LW5952, LW5953 |
| *tsp-14(jj368: tsp-14b(EQCLL-AQCAA)) sma-9(cc604); ccIs4438* | LW6020 |
| *tsp-12(jj300); tsp-14(jj368: tsp-14b(EQCLL-AQCAA)) sma-9(cc604); ccIs4438* | LW6045, LW6049 |
| TSP-14A subcellular localization | |
| *tsp-14(jj183: gfp::3xFLAG::tsp-14a); pwIs1149[snx-1p::TagRFP::rab-11]* | LW6028 |
| *tsp-14(jj183: gfp::3xFLAG::tsp-14a); pwIs1058[snx-1p::TagRFP::eea-1]* | LW6029 |
| *tsp-14(jj183: gfp::3xFLAG::tsp-14a); pwIs1116[snx-1p::TagRFP::rab-7]* | LW6030, LW6040 |
| TSP-12 and TSP-14A or TSP-14B co-localization | |
| *tsp-12(jj195: TagRFP::3xMyc::tsp-12); tsp-14(jj192: gfp::3xFLAG::tsp-14b)* | LW4634, LW4635 |
| *tsp-12(jj195: TagRFP::3xMyc::tsp-12); tsp-14(jj184: gfp::3xFLAG::tsp-14a)* | LW4654, LW4655 |
| *tsp-12(jj181: gfp::3xFlag::tsp-12); tsp-14(jj200: TagRFP::3xMyc::tsp-14a)* | LW4874 |
| Extra-chromosomal array rescue experiments | |
| *nT1[qIs51]/tsp-12(ok239); tsp-14(jj95) sma-9(cc604); arIs37(Secreted CC::gfp); jjEx4907[3.3kb Ptsp-14::tsp-14b cDNA+myo-2p::rfp]* | LW4907 |
| *nT1[qIs51]/tsp-12(ok239); tsp-14(jj95) sma-9(cc604); arIs37(Secreted CC::gfp); jjEx4912[3.3kb Ptsp-14::tsp-14b cDNA+myo-2p::rfp]* | LW4912 |
| *nT1[qIs51]/tsp-12(ok239); tsp-14(jj95) sma-9(cc604); arIs37(Secreted CC::gfp); jjEx4848[3.3kb Ptsp-14::tsp-14a cDNA, mec-7p::rfp]* | LW4848 |
| *nT1[qIs51]/tsp-12(ok239); tsp-14(jj95) sma-9(cc604); arIs37(Secreted CC::gfp); jjEx4875[3.3kb Ptsp-14::tsp-14a cDNA, mec-7p::rfp]* | LW4875 |
| *nT1[qIs51]/tsp-12(ok239); tsp-14(jj95) sma-9(cc604); arIs37(Secreted CC::gfp); jjEx4911[3.3kb Ptsp-14::tsp-14a cDNA, mec-7p::rfp]* | LW4911 |
| MosSCI insertion lines | |
| *jjSi387[3.3kb tsp-14p::tsp-14a cDNA::tsp-14 3'UTR] I; tsp-14(jj95)* | LW6224 |
| *jjSi388[3.3kb tsp-14p::tsp-14a cDNA::tsp-14 3'UTR] I; tsp-14(jj95)* | LW6230 |
| *jjSi389[3.3kb tsp-14p::tsp-14b cDNA::tsp-14 3'UTR] I; tsp-14(jj95)* | LW6225 |
| *jjSi390[3.3kb tsp-14p::tsp-14b cDNA::tsp-14 3'UTR] I; tsp-14(jj95)* | LW6226 |
| *jjSi401[5.2kb tsp-14p::tsp-14a cDNA::tsp-14 3'UTR] I; tsp-14(jj95)* | LW6244 |
| *jjSi402[5.2kb tsp-14p::tsp-14b cDNA::tsp-14 3'UTR] I; tsp-14(jj95)* | LW6245 |
| *jjSi393[snx-1p::tsp-14a cDNA-gDNA chimera::gfp::3xflag::tbb-2 3'UTR] I; tsp-14(jj95)* | LW6231 |
| *jjSi394[snx-1p::tsp-14a cDNA-gDNA chimera::gfp::3xflag::tbb-2 3'UTR] I; tsp-14(jj95)* | LW6232 |
| *jjSi395[snx-1p::tsp-14b cDNA-gDNA chimera::gfp::3xflag::tbb-2 3'UTR] I; tsp-14(jj95)* | LW6233 |
| *jjSi396[snx-1p::tsp-14b cDNA-gDNA chimera::gfp::3xflag::tbb-2 3'UTR] I; tsp-14(jj95)* | LW6234 |

| | |
|---|---|
| Functionality of MosSci lines | |
| *jjSi388[3.3kb Ptsp-14::tsp-14a cDNA::tsp-14 3'UTR]; nT1[qIs51]/tsp-12(jj300); tsp-14(jj95)* | LW6253, LW6254 |
| *jjSi390[3.3kb Ptsp-14::tsp-14b cDNA::tsp-14 3'UTR]; nT1[qIs51]/tsp-12(jj300); tsp-14(jj95)* | LW6255, LW6256 |
| *jjSi401[5.2kb Ptsp-14::tsp-14a cDNA::tsp-14 3'UTR]; nT1[qIs51]/tsp-12(jj300); tsp-14(jj95)* | LW6257, LW6258 |
| *jjSi402[5.2kb Ptsp-14::tsp-14b cDNA::tsp-14 3'UTR]; nT1[qIs51]/tsp-12(jj300); tsp-14(jj95)* | LW6259, LW6260 |
| *jjSi393[snx-1p::tsp-14a cDNA-gDNA chimera::gfp::3xflag::tbb-2 3'UTR]; nT1[qIs51]/tsp-12(jj300); tsp-14(jj95)* | LW6266, LW6267 |
| *jjSi395[snx-1p::tsp-14b cDNA-gDNA chimera::gfp::3xflag::tbb-2 3'UTR]; nT1[qIs51]/tsp-12(jj300); tsp-14(jj95)* | LW6261, LW6262 |
| *jjSi388[3.3kb Ptsp-14::tsp-14a cDNA::tsp-14 3'UTR]; nT1[qIs51]/tsp-12(jj300); tsp-14(jj95) sma-9(cc604); ccIs4438* | LW6249, LW6250 |
| *jjSi390[3.3kb Ptsp-14::tsp-14b cDNA::tsp-14 3'UTR]; nT1[qIs51]/tsp-12(jj300); tsp-14(jj95) sma-9(cc604); ccIs4438* | LW6251, LW6252 |
| *jjSi401[5.2kb Ptsp-14::tsp-14a cDNA::tsp-14 3'UTR]; nT1[qIs51]/tsp-12(jj300); tsp-14(jj95) sma-9(cc604); ccIs4438* | LW6263, LW6274 |
| *jjSi402[5.2kb Ptsp-14::tsp-14b cDNA::tsp-14 3'UTR]; nT1[qIs51]/tsp-12(jj300); tsp-14(jj95) sma-9(cc604); ccIs4438* | LW6264, LW6265 |
| *jjSi393[snx-1p::tsp-14a cDNA-gDNA chimera::gfp::3xflag::tbb-2 3'UTR]; nT1[qIs51]/tsp-12(jj300); tsp-14(jj95) sma-9(cc604); ccIs4438* | LW6272, LW6273 |
| *jjSi395[snx-1p::tsp-14b cDNA-gDNA chimera::gfp::3xflag::tbb-2 3'UTR]; nT1[qIs51]/tsp-12(jj300); tsp-14(jj95) sma-9(cc604); ccIs4438* | LW6268, LW6269 |
| Transcriptional reporters | |
| *jjEx5717[5.2kb tsp-14p::4xNLS::gfp::unc-54 3'UTR + pRF4]* | LW5717 |
| *jjEx5718[5.2kb tsp-14p::4xNLS::gfp::unc-54 3'UTR + pRF4]* | LW5718 |
| *jjEx5719[5.2kb tsp-14p::4xNLS::gfp::unc-54 3'UTR + pRF4]* | LW5719 |
| *jjEx5720[5.2kb tsp-14p::4xNLS::gfp::unc-54 3'UTR + pRF4]* | LW5720 |
| *jjEx5721[5.2kb tsp-14p::4xNLS::gfp::unc-54 3'UTR + pRF4]* | LW5721 |
| *jjEx5722[5.2kb tsp-14p::4xNLS::gfp::unc-54 3'UTR + pRF4]* | LW5722 |
| *jjEx5723[5.2kb tsp-14p::4xNLS::gfp::unc-54 3'UTR + pRF4]* | LW5723 |
| *jjEx5724[3.3kb tsp-14p::4xNLS::gfp::unc-54 3'UTR + pRF4]* | LW5724 |

**Table S2. Oligonucleotides and plasmids used in this study.**

| **Oligonucleotides for genotyping mutants and knock-ins** |
|---|
| *tsp-12(ok239)* genotyping:<br>JKL1223 (TGTGCTGCCATGTGGCTTTC, F)<br>LW47 (GTTTCGAGCAGTTTTACGCCACC, R)<br>JKL1399 (CGACGATTGGGATAGAAACACCTATTTTTC, F)<br>or<br>LW47 (GTTTCGAGCAGTTTTACGCCACC, R)<br>LW49 (GGAATCAACGGAGCCGACGAT, R)<br>LW50 (AAGTTTCGGCAGACATCCTTCCG, F) |
| *tsp-12(jj300)* genotyping:<br>ZL670 (CGTCGGCCAGAACATCATC, F)<br>ZL671 (TTTCCGATTCACCACCAGC, R)<br>ZL672 (AAACGAACACCGTGCCC, F)<br>or<br>ZL374 (CCCTCTGCTGTGCTCTGTGTTGC, F)<br>ZL375 (CTGAAGGGGAACATTTCTTTCCTGG, R)<br>ZL539 (TTTTGTGACCGGCATAGGCTGTG, F) |
| *tsp-14(jj95, jj96)* genotyping:<br>ZL178 (CGCTTGTGACTGGGAACA, R)<br>ZL179 (GACACACCGAGATACACTGAAA, F)<br>ZL180 (CAGAAGGACACGCGCTTTAT, F) |
| *tsp-14a(jj304, jj323, jj324, 325)* genotyping:<br>ZL518 (GCCGTCACTCATAACACCCATTC, F)<br>ZL675 (GACTGGGAGCAATGCCTACTAATAG, F)<br>ZL676 (GGGAATTTGCAACTTTGACC, R)<br>followed by sequencing using ZL518 |
| *tsp-14b(jj301, jj316, jj317, 318)* genotyping:<br>ZL635 (TTCCTGAAGCTAGGCAGC, F)<br>ZL636 (CATCTTGGTCCACGAACACC, R)<br>ZL641 (CCACGGCGATTCTTAAAAAC, F)<br>followed by sequencing using ZL635 |
| *tsp-14b(EQCLL-AQCAA, jj368)* genotyping:<br>ZL693 (CGCGGTTCACTGATTTCGC, F)<br>ZL694 (CTGTGGCGGCTTTGTTTGG, R)<br>ZL723 (GAGCCTGCGATTGGGC, F)<br>followed by sequencing using ZL694 or<br>ZL518 (GCCGTCACTCATAACACCCATTC, F)<br>ZL519 (CGTGGTTTGCGTTGCTTGTGTG, R)<br>ZL588 (CTGCGATTGGGCCCAATGCGCTG, F)<br>followed by sequencing using ZL518 |
| *gfp::3×flag::tsp-14b(jj192, jj193, 202, jj326)* genotyping:<br>ZL349 (CGCCGGAATCACCCACGGAATGG, F)<br>ZL428 (CGTCTTCATAAAACGTCGCGGTG, R)<br>ZL537 (CGACTTCCTGAAGCTAGGCAGC, F) |

| |
|---|
| *tsp-14::gfp::3×flag(jj219, jj319, jj327, jj377, jj378)* genotyping:<br>ZL349 (CGCCGGAATCACCCACGGAATGG, F)<br>ZL502 (CGCGTTCGAGACAGAAAGATTTGG, F)<br>ZL513 (GCGACAACGACTCGAACACCTCC, R) |
| *gfp::3×flag::tsp-12(jj181)* genotyping:<br>ZL431 (GGAGAATCTGTACTTTCAATCCGG, F)<br>ZL374 (CCCTCTGCTGTGCTCTGTGTTGC, R)<br>ZL375 (CTGAAGGGGAACATTTCTTTCCTGG, F) |
| *gfp::3×flag::tsp-14a(jj183, jj184, jj186)* genotyping:<br>ZL516 (CAAGGAGAATCTGTACTTTCAATCCGG, F)<br>ZL376 (CGCGTCGGCTCGCCGGAGCC, R)<br>ZL377 (CGTTATCCTGTTGGTCACGCAGG, F) |
| *tagRFP::3×Myc::tsp-14a(jj200, jj201)* genotyping:<br>ZL430 (TTATACGAAGTTATTTTCAGGGAGCCGG, F)<br>ZL376 (CGCGTCGGCTCGCCGGAGCC, R)<br>ZL377 (CGTTATCCTGTTGGTCACGCAGG, F) |
| *sma-9(cc604)* sequencing:<br>MLF69 (CGCAACAAGTTCATTCTCCA, F)<br>MLF70 (CTTGGCTAAGATCCCATGCT, R)<br>followed by sequencing using MLF69 |
| *tagRFP::3×Myc::tsp-12(jj195)* genotyping:<br>ZL430 (TTATACGAAGTTATTTTCAGGGAGCCGG, F)<br>ZL374 (CCCTCTGCTGTGCTCTGTGTTGC, F)<br>ZL375 (CTGAAGGGGAACATTTCTTTCCTGG, R) |

**Oligonucleotides for sgRNA plasmid construction and/or repair template used in CRISPR/Cas9-mediated genome editing.**

| |
|---|
| For pZL58 (*tsp-12* knockout #1 sgRNA):<br>ZL255 (TCTTGACTCGCTCAACGGGCAAAA)<br>ZL256 (AAACTTTTGCCCGTTGAGCGAGTC) |
| For pZL79 (*tsp-12* knockout #2 sgRNA):<br>ZL334 (TCTTGATGGCCAATCGACGACAGC)<br>ZL335 (AAACGCTGTCGTCGATTGGCCATC) |
| For pZL8 (*tsp-14* knockout #1 sgRNA):<br>LW1 (CGGGAATTCCTCCAAGAACTCGTACAAAAATGCTCT)<br>ZL28 (ACGCGAGCTCGCGCGGTGGTCAAACATTTAGATTTGCAATTCAATTATATAG)<br>LW2 (CGGAAGCTTCACAGCCGACTATGTTTGGCGT)<br>ZL27 (GACCACCGCGCGAGCTCGCGTGTTTTAGAGCTAGAAATAGCAAGTTA) |
| For pZL 9(*tsp-14* knockout #2 sgRNA):<br>LW1 (CGGGAATTCCTCCAAGAACTCGTACAAAAATGCTCT)<br>ZL30 (CGGATGCCTTCAGCCGCTTCAAACATTTAGATTTGCAATTCAATTATATAG)<br>LW2 (CGGAAGCTTCACAGCCGACTATGTTTGGCGT)<br>ZL27 (GAAGCGGCTGAAGGCATCCGGTTTTAGAGCTAGAAATAGCAAGTTA) |
| For pZL80 (*tsp-14a* specific knockout #1 sgRNA):<br>ZL336 (TCTTGGGGAACAGTGTCTGCTGAT)<br>ZL337 (AAACATCAGCAGACACTGTTCCCC) |
| *tsp-14a* specific knockout repair oligo |

| |
|---|
| ZL639 (CACATTCTTTCAGCGCACCGGGCGCTTGTGACTGGGAGCAATGC<br>CTACTAATAGGCTCCGGCGAGCCGACGCGAGCTCGCGCGGTGGTCTC) |
| For pZL83 (*tsp-14b* specific knockout #2 sgRNA):<br>ZL342 (TCTTGGCCCCATCGCGCACCGCGA)<br>ZL343 (AAACTCGCGGTGCGCGATGGGGCC) |
| *tsp-14b* specific knockout repair oligo<br>ZL638 (TTTTGTCTGAAACCATTCTGAAATAATCTACATGAGTTGCCACA<br>CCGAGCTCCACGGCGATTCTTAAAAACGTAAGTGTTGTTTTGTA<br>GAGGGAAAAGAGACACAC) |
| For pZL59 (sgRNA, C-terminally tagged TSP-14 with *jj304*, *jj317* or *jj368* mutations.)<br>ZL257 (TCTTGGTACTGCAGCAAACCGATT)<br>Zl258 (AAACAATCGGTTTGCTGCAGTACC) |
| For pZL66 (Repair template, C-terminally tagged TSP-14 with *jj304*, *jj317* or *jj368* mutations.)<br>ZL291 (ACGTTGTAAAACGACGGCCAGTCGCCGGCAATTAAATCTGCCCG<br>CTCCTCCTG)<br>ZL292 (CATCGATGCTCCTGAGGCTCCCGATGCTCCGGACTTGGACTTCTG<br>TGGGACGAGATCAGTTTGCTGCAGTACCCATTGATGC)<br>ZL293 (CGTGATTACAAGGATGACGATGACAAGAGATAAATGTTATAGTAA<br>ATGATATTCAAATTTAAC)<br>ZL294 (GGAAACAGCTATGACCATGTTATCGATTTCGTTAAACGGATCAAG<br>CAAGCAAAATC) |
| For pZL80 (sgRNA, *tsp-14b* sorting signal mutated (EQCLL – AQCAA))<br>ZL336 (TCTTGGGGAACAGTGTCTGCTGAT)<br>ZL337 (AAACATCAGCAGACACTGTTCCCC) |
| For pZL141 (Repair template, *tsp-14b* sorting signal mutated (EQCLL – AQCAA))<br>ZL585 (TCTTGCGCACCGGGCGCTTGTGAC)<br>ZL586 (AAACGTCACAAGCGCCCGGTGCGC) |
| For pZL60 (*gfp::3×flag::tsp-12* or *tagRFP::3×Myc::tsp-12* knock-in #1 sgRNA):<br>ZL267 (TCTTGCCACTTCTCACCCATCAGA)<br>ZL268 (AAACTCTGATGGGTGAGAAGTGGC) |
| For pZL79 (*gfp::3×flag::tsp-12* or *tagRFP::3×Myc::tsp-12* knock-in #2 sgRNA):<br>ZL334(TCTTGATGGCCAATCGACGACAGC)<br>ZL335(AAACGCTGTCGTCGATTGGCCATC) |
| For pZL64 (*gfp::3×flag*::*tsp-12* knock-in repair template):<br>ZL283 (ACGTTGTAAAACGACGGCCAGTCGCCGGCAACCTCCCGTCGCTT<br>TGTTCA)<br>ZL284 (TCCAGTGAACAATTCTTCTCCTTTACTCATCTGATGGGTGAGAAGT<br>GGCCGA)<br>ZL285 (CGTGATTACAAGGATGACGATGACAAGAGAATGGCCAATCGACGA<br>CAGCCAGTGCAACACAGAGCACAGCAGAG)<br>ZL286 (TCACACAGGAAACAGCTATGACCATGTTATGCTGAGAATTCGGCGA<br>TGAGC) |
| For pZL134 (*tagRFP::3×Myc*::*tsp-12* knock-in repair template):<br>ZL283 (ACGTTGTAAAACGACGGCCAGTCGCCGGCAACCTCCCGTCGCTT<br>TGTTCA)<br>ZL286 (TCACACAGGAAACAGCTATGACCATGTTATGCTGAGAATTCGGCGA |

| |
|---|
| TGAGC)<br>ZL358 (CTTGATGAGCTCCTCTCCCTTGGAGACCATCTGATGGGTGAGAAGT GGCCGA)<br>ZL359 (GAGCAGAAGTTGATCAGCGAGGAAGACTTGATGGCCAATCGACGA CAGCCAGTGCAACACAGAGCACAGCAGAG) |
| For pZL80 (*gfp::3×flag::tsp-14a* or *tagRFP::3×Myc::tsp-14a* knock-in sgRNA):<br>ZL336 (TCTTGGGGAACAGTGTCTGCTGAT)<br>ZL337 (AAACATCAGCAGACACTGTTCCCC) |
| For pZL81 (*gfp::3×flag::tsp-14a* or *tagRFP::3×Myc::tsp-14a* knock-in repair template):<br>ZL338 (ACGTTGTAAAACGACGGCCAGTCGCCGGCAACTGAGG GTGCTTGAGAGTGGC)<br>ZL339 (TCCAGTGAACAATTCTTCTCCTTTACTCATCAGCAGACAC TGTTCCCAGTCAC)<br>ZL340 (CGTGATTACAAGGATGACGATGACAAGAGAATGGGCTCC GGCGAGCC)<br>ZL341 (TCACACAGGAAACAGCTATGACCATGTTATACTGTACCGTGA CTCGGCGC) |
| For pZL83 (*gfp::3×flag::tsp-14a* or *tagRFP::3×Myc::tsp-14a* knock-in sgRNA):<br>ZL342 (TCTTGGCCCCATCGCGCACCGCGA)<br>ZL343 (AAACTCGCGGTGCGCGATGGGGCC) |
| For pZL84 (*gfp::3×flag::tsp-14a* or *tagRFP::3×Myc::tsp-14a* knock-in repair template):<br>ZL344 (ACGTTGTAAAACGACGGCCAGTCGCCGGCACCGTTTGCCTAAAT TAGATTTCGCCAC)<br>ZL345 (TCCAGTGAACAATTCTTCTCCTTTACTCATCTCATGTAGATTATTT CAGAATGGTTTCAG)<br>ZL346 (CGTGATTACAAGGATGACGATGACAAGAGAATGCCCCATCGCGCA CCGCGACGTTTTATGAAGACGTAAGTGTTGTTTTGTAGAGG)<br>ZL347 (TCACACAGGAAACAGCTATGACCATGTTATGTTGCATGTTTGAAACCC TCAAATGTTG) |
| **Oligonucleotides for sgRNA plasmid construction and/or repair template used in CRISPR/Cas9-mediated Mos single copy insertion (MosSCI) with chromosome I ttTi4348.** |
| For pZL170(*3.3kb tsp-14p::tsp-14a cDNA::tsp-14 3'UTR*, MosSCI repair template):<br>ZL735 (tgtaaaacgacggccagtgcGTAAGTGTTGTTTTGTAGAG)<br>ZL736 (ggaaacagctatgaccatgcGACAGTTTAAAACAGTAAATTTCAG) |
| For pZL171(*3.3kb tsp-14p::tsp-14b cDNA::tsp-14 3'UTR*, MosSCI repair template):<br>ZL735 (tgtaaaacgacggccagtgcGTAAGTGTTGTTTTGTAGAG)<br>ZL736 (ggaaacagctatgaccatgcGACAGTTTAAAACAGTAAATTTCAG) |
| For pZL172(*5.2kb tsp-14p::tsp-14a cDNA::tsp-14 3'UTR*, MosSCI repair template):<br>ZL739 (tgtaaaacgacggccagtgcACTCGAGAAATTAGTTGC)<br>ZL740 (caacacttacGTCTTCATAAACCGTCGC)<br>ZL741 (ttatgaagacGTAAGTGTTGTTTTGTAGAG)<br>ZL736 (ggaaacagctatgaccatgcGACAGTTTAAAACAGTAAATTTCAG) |
| For pZL173(*5.2kb tsp-14p::tsp-14b cDNA::tsp-14 3'UTR*, MosSCI repair template):<br>ZL739 (tgtaaaacgacggccagtgcACTCGAGAAATTAGTTGC)<br>ZL740 (caacacttacGTCTTCATAAACCGTCGC)<br>ZL741 (ttatgaagacGTAAGTGTTGTTTTGTAGAG) |

ZL736 (ggaaacagctatgaccatgcGACAGTTTAAAACAGTAAATTTCAG)

For pJKL1226(*snx-1p::tsp-14a cDNA-gDNA chimera::gfp::3xflag::tbb-2 3'UTR*, MosSCI repair template):

JKL1940 (TCGATCATCCtgtaaaacgacggccagtgcGAAGGAATTGTCGAGCTTTTCA GTTTCTTTTAC)
JKL1938 (GTCGGCTCGCCGGAGCCCATCCGGTTACCTTCCAGGTGGAC)
JKL1939 (GTCCACCTGGAAGGTAACCGGATGGGCTCCGGCGAGCCGAC)
JKL1941 (gacactttttgggagttcaaggcttacACATTTGAGCAAGAAAATATTGTCGC GCAAG)
JKL1942 (CTTGCGCGACAATATTTTCTTGCTCAAATGTgtaagccttgaactcccaaaaagtgtc)
JKL1943 (tgcttgaaaggattttgcatttatcTTAAGTTATCTCTTGTCATCGTCATC CTTGTAATC)

For pJKL1227(*snx-1p::tsp-14b cDNA-gDNA chimera::gfp::3xflag::tbb-2 3'UTR*, MosSCI repair template):

JKL1940 (TCGATCATCCtgtaaaacgacggccagtgcGAAGGAATTGTCGAGCTTTTCA GTTTCTTTTAC)
JKL1944 (CGCGGTGCGCGATGGGGCATCCGGTTACCTTCCAGGTGGAC)
JKL1945 (GTCCACCTGGAAGGTAACCGGATGCCCCATCGCGCACCGCG)
JKL1941 (gacactttttgggagttcaaggcttacACATTTGAGCAAGAAAATATTGTCGC GCAAG)
JKL1942 (CTTGCGCGACAATATTTTCTTGCTCAAATGTgtaagccttgaactcccaaaaagtgtc)
JKL1943 (tgcttgaaaggattttgcatttatcTTAAGTTATCTCTTGTCATCGTCATC CTTGTAATC)

**Primers used to confirm genomic DNA sequence for knock-ins and knockouts**

*tsp-12(jj300)* sequencing:

ZL670 (CGTCGGCCAGAACATCATC)
ZL672 (AAACGAACACCGTGCCC)
followed by sequencing using ZL670

*tsp-12(jj300)* sequencing:

ZL375 (CTGAAGGGGAACATTTCTTTCCTGG, R)
ZL539 (TTTTGTGACCGGCATAGGCTGTG, F)
followed by sequencing using ZL539

*tagRFP::3×Myc::tsp-12(jj257, jj258)*

ZL531(CTGATTTCCGATTCACCACCAGCG, F)
ZL375 (CTGAAGGGGAACATTTCTTTCCTGG, R)
followed by sequencing using ZL531

*tsp-14::gfp::3×flag(jj218, jj219)*

ZL536 (GGAAAGAGAGATGGGGTGGGG, F)
ZL513 (GCGACAACGACTCGAACACCTCC, R)
followed by sequencing using ZL536

*tsp-14::tagRFP::3×Myc(jj265, jj266)*

ZL513 (GCGACAACGACTCGAACACCTCC, R)
ZL536 (GGAAAGAGAGATGGGGTGGGG, F)
followed by sequencing using ZL536

*tsp-14a(jj304, jj323, jj324, 325)* sequencing:

ZL693 (CGCGGTTCACTGATTTCGC, F)

| |
|---|
| ZL694 (CTGTGGCGGCTTTGTTTGG, R)<br>followed by sequencing using ZL693 |
| *tsp-14b(jj301, jj316, jj317, 318)* sequencing:<br>ZL691 (GCCACCCTCATCCACTGC, F)<br>ZL692 (CAATTACAATTTTCTGTGGGGTTCG, R)<br>followed by sequencing using ZL691 |
| **Primers used to confirm genotype of different MosSCI knock-ins** |
| *jjSi388, jjSi390, jjSi401, jjSi402* genotyping:<br>ZL746 (GCAGAAATACCTCCCTGTCAATTCC)<br>ZL747 (TGAGCGTATCTATCAAGTCCTTGTCTC)<br>ZL748 (ACGGATTTAGTGACAGTTTCTTGGG) |
| *jjSi393, jjSi395* genotyping:<br>ZL746 (GCAGAAATACCTCCCTGTCAATTCC)<br>ZL747 (TGAGCGTATCTATCAAGTCCTTGTCTC)<br>ZL763 (GACGTGGGGGGAATTGGTGG) |
| **Primers used to confirm genomic DNA of different MosSCI knock-ins** |
| *jjSi388, jjSi390, jjSi393, jjSi395, jjSi401, jjSi402* genomic DNA sequencing:<br>ZL749 (GCGGAATACGAATTGGGAGACG)<br>ZL750 (TGAAATCTGAAGCACTGCCGC)<br>Sequencing with both ZL749 and ZL750.<br>ZL377 (CGTTATCCTGTTGGTCACGCAGG)<br>ZL751 (GCGATGGCTAGAATTCCGGAC)<br>Sequencing with both ZL377 and ZL751.<br>ZL752 (GTCCGGAATTCTAGCCATCGC)<br>ZL747 (TGAGCGTATCTATCAAGTCCTTGTCTC)<br>Sequencing with both ZL752 and ZL747. |
